# Directed evolution of the *Drosophila* microbiome improves intestinal health and extends lifespan

**DOI:** 10.64898/2026.08.04.742805

**Authors:** Matt Ulgherait, Yiwei Sun, Yiming Huang, Timothy Chang, Carly Lam, Julie Canman, Harris Wang, Mimi Shirasu-Hiza

## Abstract

The gut microbiome and its bacterially derived metabolites are known to affect many aspects of the host organism’s health, including metabolism, immune response, intestinal inflammation, oxidative stress, and even lifespan. Because pathological changes in the gut microbiome and these functions are associated with aging, many have hypothesized that we could protect against aging by generating beneficial changes to the gut microbiome. Here, we directed evolution outside of the host (*ex vivo*) and generated a *Drosophila* gut microbiome resistant to paraquat, a toxin that causes oxidative stress. Compared to a control microbiome, this paraquat-resistant (PQR) microbiome transplanted back into the *Drosophila* gut endowed the host with multiple health benefits: increased resistance to dietary paraquat, reduced age-related pathologies in the gut, and extended lifespan. We identified the beneficial species of the PQR microbiome as *Lactiplantibacillus plantarum* and further identified mutations specific to lifespan-extending isolates linked to greater production of acetate. Directly feeding this short-chain fatty acid, acetate, to *Drosophila* was sufficient to recapitulate an extended lifespan, similar to that induced by gut colonization of PQR bacteria in the gut. These results serve as a proof of principle that increasing the resistance of the microbiome to oxidative stress via directed *ex vivo* evolution could serve as a therapeutic strategy to protect against aging.

## INTRODUCTION

From fruit flies to humans, the gut microbiome (bacteria and other microbes residing in the intestine) is known to have significant impact on physiological functions of the host, including metabolism, immune response, intestinal inflammation, and longevity.^1–5^ All these physiological functions can contribute to aging.^6^ In particular, aging in both flies and vertebrates is correlated with increases in gut microbial load and a shift in composition to bacterial populations associated with pathological consequences.^7,8^ In *Drosophila*, aging-related changes in gut microbiota also induce specific immune response proteins such as Dual Oxidase (DUOX), which mediates reactive oxygen species (ROS) production.^9,10^ Gut ROS production defends against microbial overgrowth, but excessive ROS production by DUOX also leads to pathology: damage to the host epithelium, aberrant proliferation of intestinal stem cells (ISCs) and enteroblasts (EBs), disorganization of the epithelia, and intestinal barrier dysfunction.^11–14^ Overexpression of antioxidant enzymes in ISCs both prevents age-related dysbiosis of the intestine and also extends lifespan.^11^ Thus, aging is associated with pathological changes in the gut microbiota, age-related dysbiosis, and increased ROS production.^15^

These observations raise the possibility that deliberately altering the gut microbiome could benefit the host and protect against the detrimental effects of aging.^2,16^ With its powerful genetic tools and short lifespan, *Drosophila* has become an important model organism for studying intestinal physiology, aging, and more recently, the gut microbiome.^4,17^ The molecular pathways underlying intestinal stem cell functions, innate immune response, and aging-related pathologies of the gut are all highly evolutionarily conserved.^6,17,18^ The potential benefits of altering the gut microbiota to promote beneficial bacteria, which could not only thrive in a high ROS environment but also resolve oxidative stress and alter gut physiology, remain largely unexplored.

Here we show that, through *ex vivo* directed evolution, we generated a gut microbiome capable of growth even in the presence of high levels of paraquat, a toxin that leads to excessive ROS production. This modified gut microbiome, termed paraquat-resistant (PQR), provided health benefits to the host *Drosophila* once transplanted into the fly. We found that *Drosophila* populated with the PQR microbiome had improved stress resistance, increased lifespan, and delayed markers of intestinal aging. Furthermore, we identified *Lactiplantibacillus plantarum* as the species in this modified microbiome that conferred this lifespan benefit to the host. We also identified mutations common to PQR clones in a single operon that are predicted to alter metabolite production of *L. plantarum*, resulting in increased acetate secretion. Dietary supplementation of acetate to adult flies was sufficient to extend lifespan similar to the lifespan extension seen with PQR bacteria. Taken together, our results serve as a proof of concept that changing the genetic makeup of the microbiome via directed evolution can serve as a therapeutic strategy to reduce intestinal inflammation, ROS production, and extend lifespan.

## RESULTS

### Directed evolution of the *Drosophila* microbiome against ROS extends fly lifespan

To generate an oxidative stress-resistant *Drosophila* microbiome, we used a strategy of directed *ex vivo* evolution. We cultured intestinal bacteria of wild-type *Drosophila* in MRS media over several weeks, split into two cultures: a control culture that was diluted with media over time and an experimental culture diluted with media containing gradually increasing concentrations of paraquat (a pesticide and ROS-producing electron transport chain disruptor; see Methods). After 25 passages, equivalent to hundreds of generations (**Figure 1A**), the experimental microbial population, which we call paraquat-resistant or “PQR”, was more capable of growth in high levels of paraquat (100 mM) than the control microbial population (**Figure 1B**). To test whether the PQR microbes can confer benefits to the host *Drosophila*, we colonized axenic females with control or PQR populations (**Figure 1C**). After 5 days, these flies were then fed media containing 20 mM paraquat, which resulted in the PQR-colonized flies surviving significantly longer than their control-colonized comparison (**Figure 1D**, solid lines). High doses of paraquat were sufficient to kill off the control gut microbiome, while PQR colonized flies maintained intestinal bacteria upon paraquat challenge (**Figure S1A, B**). This suggests the PQR microbiota can protect the host against an exogenous oxidative stressor such as paraquat. To ensure that this benefit is solely due to the continued presence of the PQR microbiome, we fed both control and PQR flies a large dose of antibiotics to eradicate their intestinal bacterial populations. With antibiotic treatment, both sets of flies had the same survival time after paraquat feeding, suggesting that the paraquat survival benefit was due to the PQR microbial population (**Figure 1D**, dashed lines).

**Figure 1.**
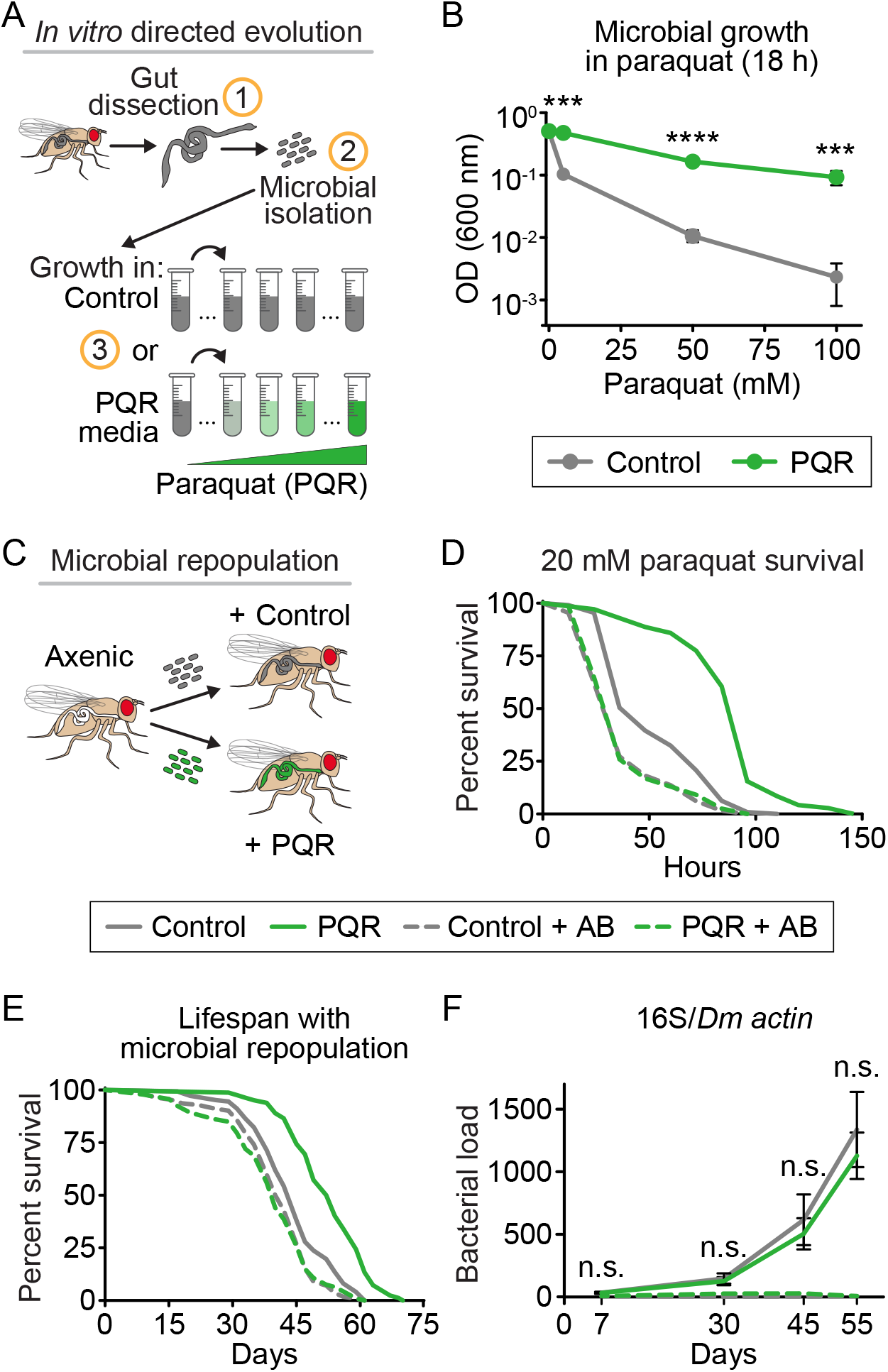
*Drosophila* colonized with *ex vivo* evolved PQR microbiome survived paraquat feeding longer and had extended lifespans relative to flies colonized with control microbiome. (**A**) Schematic of protocol for *ex vivo* evolution of the *Drosophila* microbiome in flies treated with either vehicle control (water) or paraquat (PQR), a ROS-producing toxin. (**B**) Relative to control bacteria (gray), PQR bacteria (green) grew to higher levels in media containing paraquat (***=p≤0.001, ****=p≤0.0001). (**C**) Schematic of *Drosophila* bacterial microbiome gut colonization/repopulation assay. (**D**) Relative to control-populated flies (gray solid line), PQR-populated flies (green solid line) were more resistant to paraquat toxicity (p<0.0001); this effect was ablated by antibiotic (AB) treatment to eradicate all colonized intestinal bacteria (control=gray, PQR=green, dashed lines, p>0.05). (**E**) In standard conditions (no paraquat), PQR flies (green solid) lived longer than controls (gray solid, p<0.0001); this effect was ablated by antibiotic (AB) treatment (dashed lines, p>0.05). (**F**) PQR and control flies did not have differences in bacterial load, as determined by universal 16S bacterial rDNA sanger sequencing (n.s.=p>0.05 for all time points, comparing either solid lines or dashed lines). p-values were obtained by ANOVA followed by Tukey’s multiple comparisons (B, F) and by logrank analysis (D, E). See Table S1 for statistical comparisons and population size for all experiments.

To investigate the influence of the PQR microbiome on aging and mortality, we tested the effects of the PQR microbiota on unchallenged (no paraquat) host lifespan. Female *Drosophila* repopulated with PQR microbiota lived ∼20% longer than control-colonized flies, with male flies also living longer though to lesser degree, ∼12% longer (**Figure 1E**, solid lines, **Table S1**). Flies that were reared conventionally had similar lifespan to that of control-colonized flies (**Table S1**). To confirm this benefit was due to the microbiome alone, we again showed that antibiotic-treated flies had no significant lifespan difference (**Figure 1E**, dashed lines). To determine if there were differences in bacterial colonization and growth with age, we tested microbial load over time via universal 16S bacterial rDNA sanger sequencing. We showed that the benefits of PQR microbiota on fly lifespan were not due to a difference in the number of bacteria found in the intestine (**Figure 1D**). Another potential reason for lifespan extension could be a difference in feeding behavior. Using the capillary feeding (CAFE) assay, we showed that there was no difference in feeding rate for flies 10 days post-colonization with either the control or PQR microbiome (**Figure S1C**). These results suggest the PQR bacteria confer a novel survival benefit to the host *Drosophila* possibly through modifying intestinal health with age.

### PQR-populated animals show decreased markers of age-related inflammation, ROS production, gut dysplasia, and intestinal barrier dysfunction

To elucidate the benefit of the PQR microbiome on the host *Drosophila* gut, we looked at its effect on well-established intestinal aging markers as potential mechanisms for lifespan extension. During aging, ROS production leads to many pathologies such as dysplasia and intestinal stem cell mis-differentiation.^11,12,14^ We tested if PQR *L. plantarum* microbiota lowered oxidative stress in the gut of aged animals *in vivo*. Staining of 45-day old PQR-colonized fly intestines with a superoxide-sensitive dye (DHE) showed that these flies had lower ROS levels when compared to the control-colonized intestines (**Figure 2A, B**). These flies also had fewer mis-differentiated ISC/EB clusters as marked by the intestinal stem cell label *esg*>GFP (**Figure 2A, C**). To investigate intestinal barrier dysfunction, another marker of aging, we performed the “Smurf” assay, in which a blue dye is co-ingested with food and remains in the intestine in healthy flies but leaks into the fly body if the intestinal barrier is permeable; gut barrier permeability naturally increases with age (**Figure 2D, left**).^12,19^ We found that flies populated with the PQR microbiome had decreased gut permeability over time (increased blue dye in body) when compared to flies with the control microbiome (**Figure 2D, right**). These results suggest that the PQR microbiome may have a protective effect against age-related ROS production, which may lead to improved intestinal health with age and increased lifespan.

### *Lactiplantibacillus plantarum* is the modified bacteria in the PQR consortium responsible for lifespan benefit

To investigate the composition of the PQR and control microbiomes, we assessed the bacterial species present in each limited MRS-grown population via 16S rDNA sequencing of ∼30 colonies per condition. We found that the PQR population had increased percentages of *Lactiplantibacillus brevis* and *Lactiplantibacillus plantarum* and reduced percentage of *Acetobacter pasteurianus* compared to the control population. *L. plantarum* was the most abundant species in the PQR population at 59% (**Figure 3A**). Additionally, of the species present, only *L. plantarum* was capable of sustained overnight growth in media containing high levels of paraquat (**Figure 3B**). Furthermore, axenic flies repopulated with our PQR *L. plantarum* strain had significantly increased survival time when fed media containing 20 mM paraquat, compared to axenic flies repopulated with our control *L. plantarum* (**Figure 3C**). In contrast, axenic flies repopulated with the other PQR monomicrobial populations had no survival benefit (**Figure S2A, B**). This result suggests that *L. plantarum* alone is the bacterial species responsible for the host benefit we see when repopulating *Drosophila* with our PQR microbiome.

**Figure 2.**
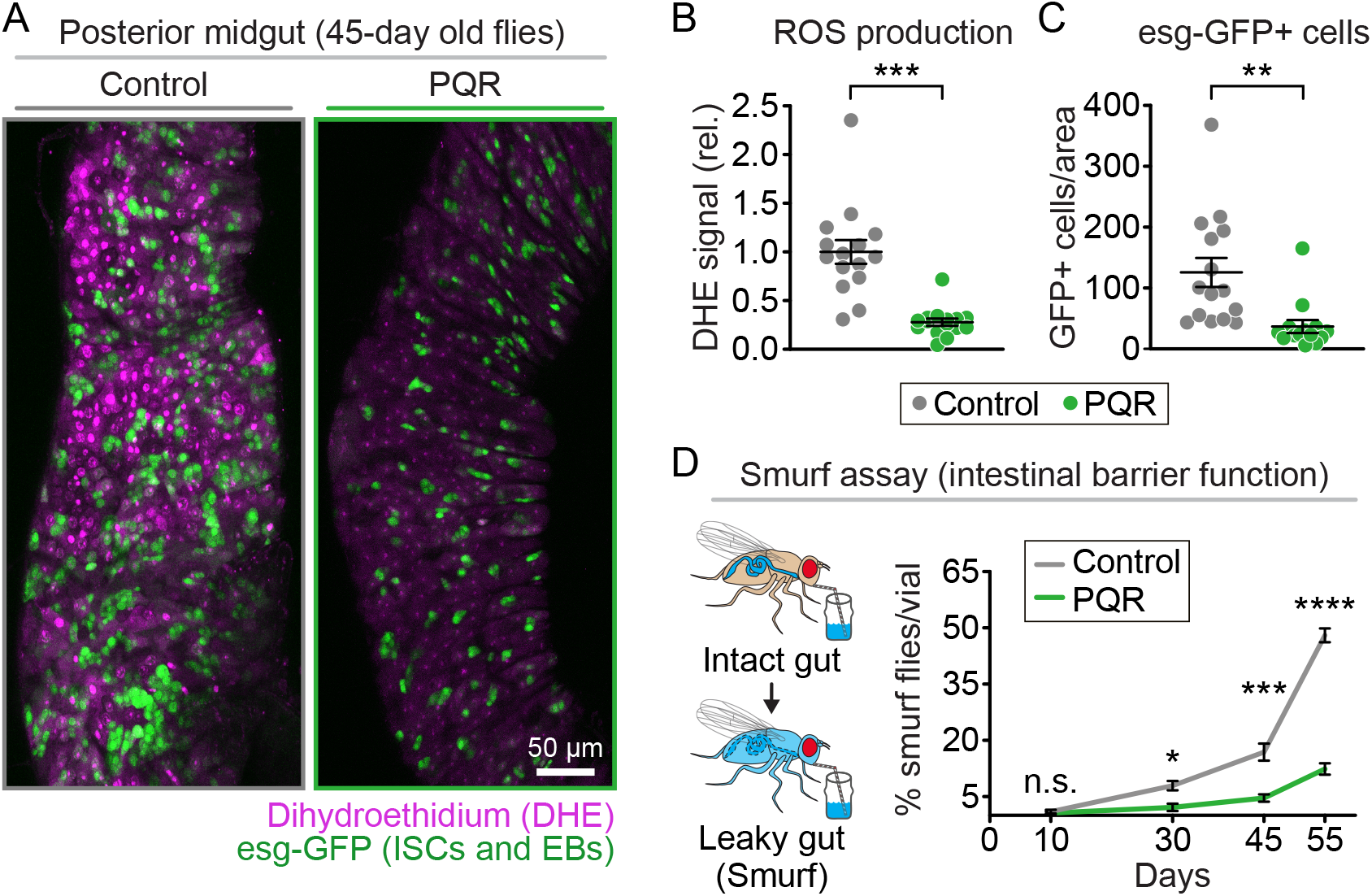
PQR microbiome colonization decreased anti-aging markers in host *Drosophila* intestine. (**A**) Representative images of guts from 45-day old flies with and without PQR microbiome colonization stained for ROS (dihydroethidium or DHE, magenta) and intestinal stem cells (ISCs) and enteroblasts (EBs), both marked by *escargot*-driven expression of GFP (*esg*>GFP, light green), scale bar=50 µm. Relative to control-populated flies (gray), PQR-populated flies (green) had (**B**) decreased DHE staining (p<0.0001) and (**C**) decreased ISC/EB numbers (p=0.0026). (**D**) (left) Schematic of the “Smurf” assay, in which flies are fed a blue dye that is normally contained within the intestine; blue dye in fly bodies indicates a loss of gut barrier function. (right) Quantitation of percent smurfing showed that PQR-populated flies (green) exhibited less smurfing than control-populated flies (gray) with increased age (p-values= n.s.:>0.05; *: ≤0.05. ***: ≤0.001; ****: ≤0.0001). p-values were obtained by 2-tailed Student’s T-test (B, C) and by ANOVA followed by Tukey’s multiple comparisons (D). See Table S1 for statistical comparisons and population size for all experiments.

**Figure 3.**
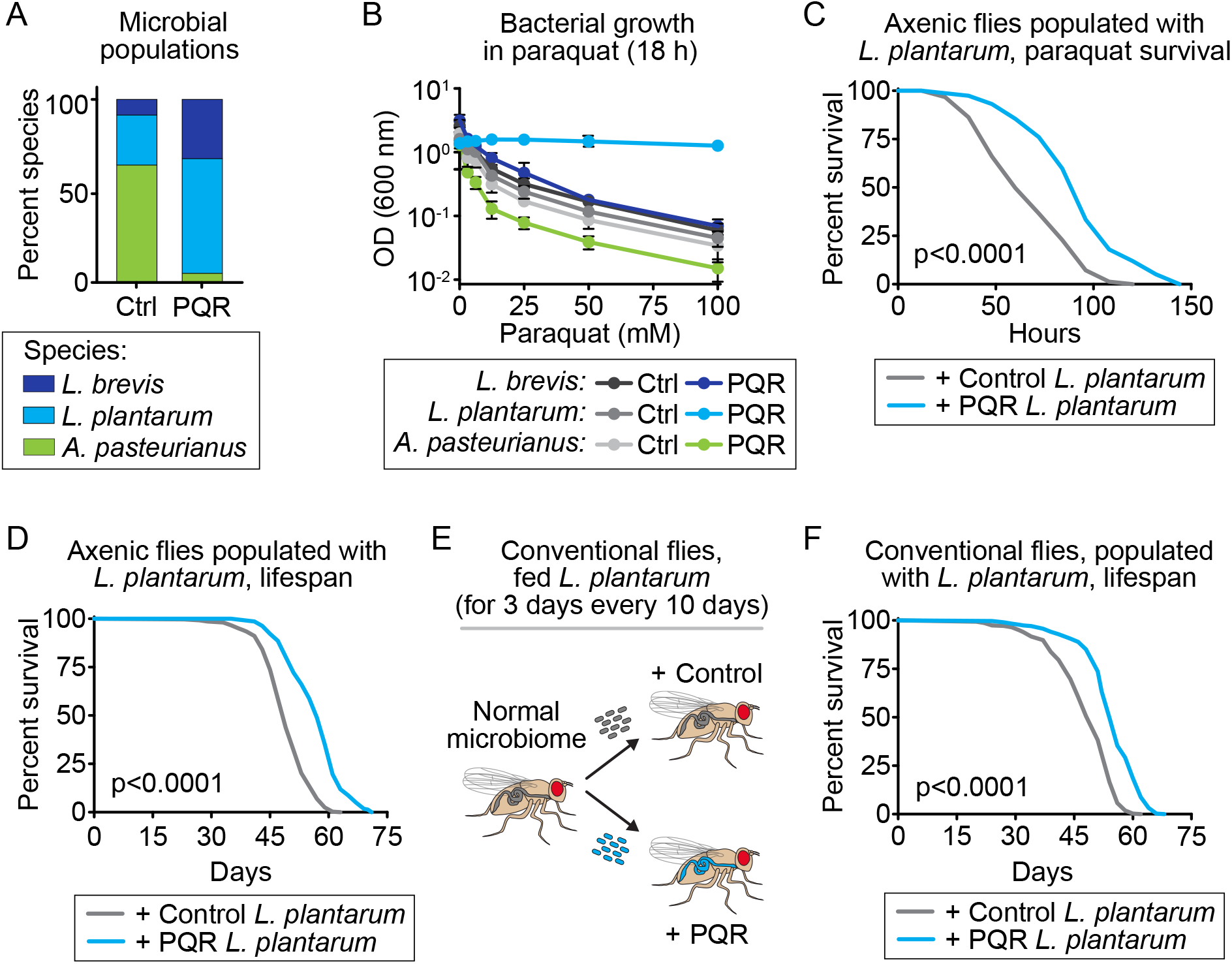
*Lactiplantibacillus plantarum* isolated from PQR evolved microbiome and transplanted into host *Drosophila* decreased anti-aging markers and increased lifespan in host. (**A**) After *ex vivo* evolution, the PQR microbiome had higher percentages of *L. brevis* (dark blue) and *L*. *plantarum* (light blue) species and a lower percentage of *A. pasteurianus* (light green) than the control microbiome, as determined by universal 16S bacterial rDNA sanger sequencing. (**B**) When single colony isolates of each species from either control microbiome (grays) or PQR microbiome (colors) were grown in media containing paraquat, PQR *L. plantarum* (light blue) grew better than *L. brevis*, *L. plantarum*, and *A. pasteurianus* (light, medium, and dark gray, respectively, p<0.0001 for all) in control media and better than *L. brevis* (dark blue, p<0.0001) and *A. pasteurianus* (green, p<0.0001) in media containing paraquat. Relative to axenic flies populated with control-treated *L. plantarum*, axenic flies populated with PQR *L. plantarum* survived longer (**C**) when challenged with paraquat (p<0.0001) and (**D**) in standard lifespan assay (p<0.0001). (**E**) Schematic of probiotic treatment of conventionally reared (not axenic) flies, fed either control (gray) or PQR (light blue) *L. plantarum* for 3 days every 10 days through their lifespan. (**F**) Probiotic treatment of conventionally reared flies with PQR *L. plantarum* (light blue) also extended lifespan relative to those treated with control *L. plantarum* (gray, p<0.0001). p-values were obtained by ANOVA followed by Tukey’s multiple comparisons (B) and by logrank analysis (C, D, F). See Table S1 for statistical comparisons and population size for all experiments.

To test whether this species can recapitulate the lifespan effect seen with the PQR microbiota, we isolated five colonies each from plated cultures of control *L. plantarum* or PQR *L. plantarum* and tested their effects on fly lifespan (**Figure 3D, Table S1**). The axenic host flies repopulated with PQR *L. plantarum* isolates had significantly longer lifespans than those repopulated with control *L. plantarum* isolates (**Figure 3D**). Thus, PQR *L. plantarum* isolate cultures were sufficient to confer a survival benefit to and extend the lifespan of the host *Drosophila*. To investigate whether this effect transfers to conventional (non-axenic) flies, we fed flies with a normal microbiome either control or PQR *L. plantarum* for 3 days every 10 days and assessed lifespan (a probiotic-like paradigm, **Figure 3E**). We found that the PQR *L. plantarum*-fed flies had an increased lifespan compared to control-fed flies (**Figure 3F**). Thus, PQR *L. plantarum* can be fed to conventional flies and confer lifespan extension, demonstrating its sufficiency and pro-biotic capability. This effect was not due to differences in bacteria number in the aging host intestine (**Figure S2C**). Taken together, these results show that PQR *L. plantarum* is responsible for the lifespan extension and survival benefit to *Drosophila*.

### Increased acetate production by gut bacteria and increased acetate in food are both associated with *Drosophila* lifespan extension

To identify genomic changes that might be responsible for PQR *L. plantarum* host benefits, including lifespan extension, we performed DNA-sequencing and SNP analysis on the five individually isolated colonies of PQR *L. plantarum*, each of which extended *Drosophila* lifespan, and compared these to the five colonies of control *L. plantarum*, none of which extended Drosophila lifespan (**Table S1**). Our analysis found that all five PQR *L. plantarum* colonies had diverse mutations in a specific operon, containing the genes *ywnA*, *pox5*, and *hmo* (**Figure 4A**), while none of the five control isolates had mutations in this operon. This result suggests the possibility that the *ywnA-pox5-hmo* operon may be responsible for PQR *L. plantarum* host benefits.

To identify corresponding changes in gene expression that might drive host benefits, we also performed RNA-sequencing analysis on two PQR isolates and two control isolates of *L. plantarum*. Each isolate was incubated in media with or without paraquat (0, 5, 20, or 50 mM paraquat) and replicate samples (a, b, or c) were taken after 0, 0.75, 2, and 10 hours of incubation for RNA preparation and sequencing. RNA seq analysis identified 66 genes differentially expressed between at least one PQR isolate and one control isolate at one time point (**Figure S3**). We found that all three genes in the *ywnA-pox5-hmo* operon were ∼50-fold overexpressed by both PQR isolates but not by control isolates, with or without paraquat, at every timepoint tested. Taken together, these results suggested that the *ywnA-pox5-hmo* operon was of interest for the paraquat resistance in *L. plantarum* and for the lifespan benefit conferred to the host *Drosophila*.

After many attempts and technical hurdles in trying to clone or even synthesize this operon or *pox5* gene, we instead turned to testing the predicted functions of this operon. One major metabolic product of this operon, due to the activity of the enzyme Pox5, is acetyl phosphate, which can readily be converted to free acetate by acetate kinase (**Figure 4C**).^20,21^ Previous studies showed that bacterially derived acetate can greatly influence *Drosophila* intestinal physiology and immune response.^22–24^ To test if PQR *L. plantarum* isolates produce more acetate than control isolates, we grew control and PQR *L. plantarum* cultures for 18 hours in MRS medium and measured acetate in the growth medium. We found that PQR *L. plantarum* cultures contained ∼5-fold higher levels of free acetate in the growth medium compared to control *L. plantarum* (**Figure 4D**). Consistent with the hypothesis that PQR *L. plantarum* provides health benefits to its host through acetate production, we found that supplementation of *Drosophila* food with sodium acetate increased lifespan of control-populated animals, but not that of PQR-populated animals (**Figure 4E**). While we observed an acetate dose-dependent increase in lifespan for flies populated with control *L. plantarum*, increasing acetate only decreased lifespan for flies populated with PQR *L. plantarum*. These data suggest that PQR bacteria produce enough acetate to facilitate lifespan extension and any additional acetate is detrimental in this context. Acetate feeding also resulted in lifespan increases in control flies with or without presence of an intestinal microbiome (**Figure S4**). These data suggest that acetate secretion by the microbiome is essential to *Drosophila* intestinal homeostasis and longevity, as suggested by other data examining microbiota metabolites.^25^ Thus, our data suggest that modification of the gut bacterium by *ex vivo* evolution against an oxidative stressor applies a selective pressure toward increasing microbial acetate production and that this increased acetate in the gut is sufficient to extend host lifespan.

**Figure 4.**
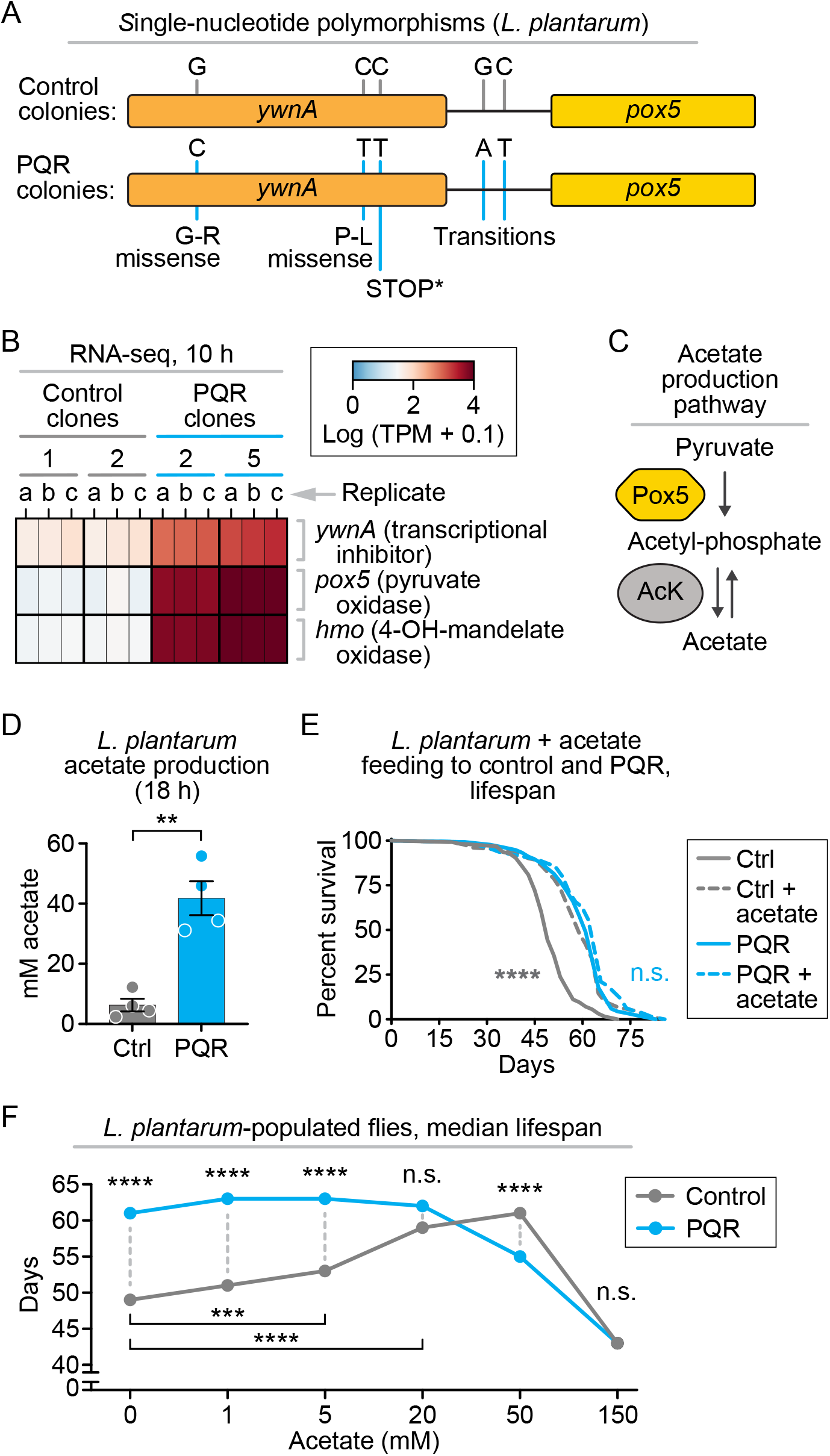
Multiple PQR *L. plantarum* clones that increased host lifespan had mutations in a common operon and exhibited increased production of acetate, which extended lifespan when added exogenously. (**A**) DNA-seq single-nucleotide polymorphism (SNP) analysis of five independent PQR-evolved *L. plantarum* clones revealed mutations in a single common operon, *ywnA*-*pox5*-*hmo*, mutations predicted to inhibit YwnA activity, with no similar mutations in five independent control *L. plantarum* isolates. (**B**) YwnA is thought to inhibit expression of its operon; RNA-seq analysis confirmed that the *ywnA*-*pox5*-*hmo* operon was over-expressed (∼50-fold) in two PQR *L. plantarum* isolates (light blue, clones 2 and 5, triplicate samples a-c) relative to two control isolates (gray, clones 1 and 2, triplicate samples a-c) after 10 hours of incubation in media without paraquat. (**C**) Schematic depicting enzymatic function of Pox5 in acetate production. (**D**) Cultures of PQR *L. plantarum* (light blue) secreted 5-fold more acetate than control *L. plantarum* cultures (gray, p=0.001). (**E**) Supplementing with dietary acetate (20 mM) extended the lifespan of flies populated with control *L. plantarum* (gray, p<0.0001) but did not further extend the lifespans of flies populated with PQR *L. plantarum* (light blue, n.s. or p>0.05). (**F**) Different concentrations of dietary acetate caused lifespan extension when fed to flies populated with control *L. plantarum*, but not when fed to flies populated with PQR *L. plantarum*. p-values were obtained by two-tailed Student’s t-test (D) and by logrank analysis (E, F). See Table S1 for statistical comparisons and population size for all experiments.

## DISCUSSION

In this study, we showed that directed *ex vivo* evolution of the *Drosophila* gut microbiome using an extracellular stressor (paraquat, a toxin known to increase ROS production) led to genetically altered bacteria that, when transplanted back into the host, provided the host with significant health benefits. These health benefits included increased survival of dietary paraquat, decreased ROS levels in the gut, and lifespan extension. We identified *L. plantarum* as the critical paraquat-adapted (PQR) microbiome species conveying these host benefits and further found PQR-specific mutations and altered expression of *L. plantarum* genes involved in acetate synthesis, resulting in high levels of acetate secretion. Finally, we showed that direct dietary supplementation of acetate alone was sufficient to recapitulate the influence of PQR *L. plantarum* on host health and lifespan. Importantly, acetate supplementation did not extend the lifespan of flies whose lifespans were already extended by PQR *L. plantarum*, suggesting a maximal lifespan extension by this mechanism.

Directed evolution of the gut microbiome to extend lifespan has been attempted before with varying success. In a recent example similar to our work, *C. elegans* fed an *E. coli* OP50 that was adapted to be resistant against paraquat also experienced extended lifespan.^26^ In that case, the authors showed that this host lifespan benefit was due to high bacterial iron content, which activated host MAPK signaling and extended lifespan. This is likely effective because *C. elegans* are typically fed this single species of bacteria (*E. coli* OP50), on which they rely for sustenance. In contrast, our paraquat-evolved *Lactiplantibacillus* species appears to extend host *Drosophila* lifespan by a completely different mechanism, through increased acetate secretion and changes in oxidative stress levels in the gut itself. We hypothesize that this difference could be due to the more diverse gut microbiome composition of *Drosophila* and its more complex relationships with host physiology.

Short-chain fatty acids (SCFAs) such as acetate, butyrate, and propionate are known to be crucial compounds produced by gut microbiota in hosts ranging from *Drosophila* to mammals. SCFAs can act as cellular fuel and as signaling messengers, modulating immunity and even gene expression.^27^ In particular, acetate is a key metabolite in the production and storage of cellular energy.^23,28^ Increased bacterially derived acetate has been associated with better metabolic outcomes including reduced obesity and insulin resistance.^29,30^ Previous work has shown that increased acetate secretion by the microbiome can help mitigate inflammation in the gut and gut barrier dysfunction.^30–33^ Exogenous acetate supplementation has also been shown to reduce inflammatory markers and Inflammatory Bowel Diseases (IBDs) pathologies.^33^ Consistent with this, our PQR microbiome produced higher acetate levels and reduced age-related markers of intestinal pathology, including oxidative stress. Acetate is also known to have many antioxidant properties.^23,34,35^ Acetate may directly reduce ROS, or help rewire host-metabolism to improve ROS-related redox stress.^23,30,31,33,34^ That said, the host benefits of bacterially derived acetate are not unlimited.^32^ In our hands, higher levels of acetate (>50 mM) resulted in toxicity and lifespan shortening in the fly (**Figure 4**). The specific mechanisms by which exogenous or bacterially derived acetate mediates host benefits such as decreased oxidative stress in the gut and lifespan extension and the potential limits of these host benefits represent an important area of exploration in host-microbe interaction.

From *Drosophila* to mammals, it has long been known that ROS plays a critical role in gut homeostasis and signaling and that the gut microbiome can significantly influence intestinal ROS levels and disease states.^11,13,14,36–38^ Excess ROS production due to microbial dysbiosis/imbalance can contribute to many IBDs and the progression of colorectal cancer.^14,39^ Indeed, modification of the microbiome population has been shown to help ameliorate the progression of these diseases.^40^ Many of these approaches involved either transplantation of microbiomes from other “healthier” donors or direct genetic modification of specific bacterial enzymes.^2,41,42^ In contrast to these approaches, here we present proof-of-principal evidence to suggest that one can perform directed *ex vivo* evolution of native host gut microbes with an exogenous stressor (oxidative stress), leveraging the bacteria’s natural ability to rapidly mutate, to generate adapted bacterial strains that can provide specific health benefits once transplanted back into the host intestinal environment. In the future, this could provide an effective and unbiased strategy for microbiome modification for therapeutic intervention in intestinal disease or even aging.

---

**Figure S1.**
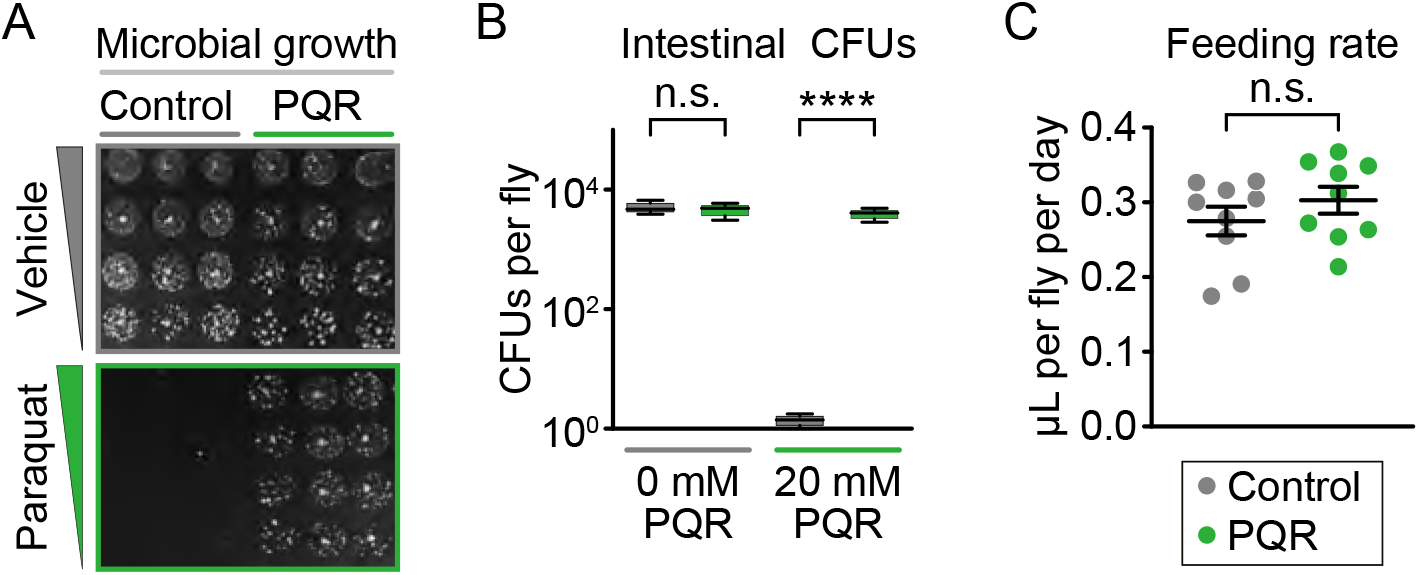
PQR microbiome recovered from the host survived paraquat media better than control microbiomes; PQR-populated flies did not exhibit significant differences in feeding rates. (**A**) Representative colony-forming units (CFU) assay for extracts from flies populated with either control (gray) or PQR (green) microbiomes plated on MRS medium with and without paraquat (20 mM). *Drosophila* showed no difference in bacterial load on media without paraquat (p>0.05) but only PQR bacteria (green) were capable of growth on medium containing paraquat (p<0.0001). (**B**) In capillary feeding (CAFE) assays, axenic *Drosophila* populated with either control (gray) or PQR-evolved (green) microbiomes showed no difference in feeding rate (n=9 each, p>0.05). p-values were obtained by ANOVA followed by Tukey’s multiple comparisons (A) or two-tailed t-test (B). See Table S1 for statistical comparisons and population size for all experiments.

**Figure S2.**
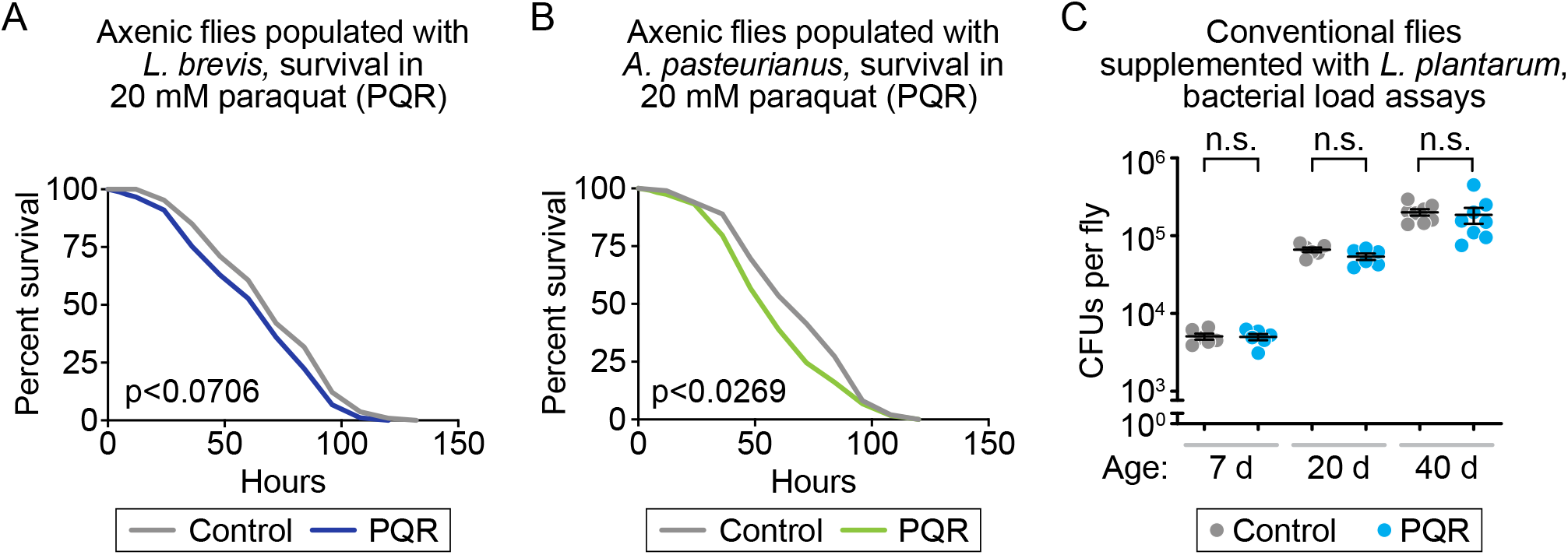
PQR-evolved *A. pastuerianus* or *L. brevis* did not confer paraquat resistance to axenic hosts, and conventional flies populated with PQR-evolved *L. plantarum* had similar bacterial loads to flies populated with control *L. plantarum* at different ages. Flies survived similarly upon paraquat challenge when populated with (**A**) control (gray) or PQR (dark blue) *L. brevis* (p=0.0706); or (**B**) control (gray) or PQR (light green) *A. pasteurianus* (p=0.0269). (**C**) Conventionally reared (conventional) flies having microbiomes supplemented with either control (gray) or PQR-evolved (light blue) *L. plantarum* had similar bacterial loads over time (p>0.05 for control vs. PQR comparisons). p-values obtained by logrank analysis (A, B) and ANOVA followed by Tukey’s multiple comparisons (C). See Table S1 for statistical comparisons and population size for all experiments.

**Figure S3.**
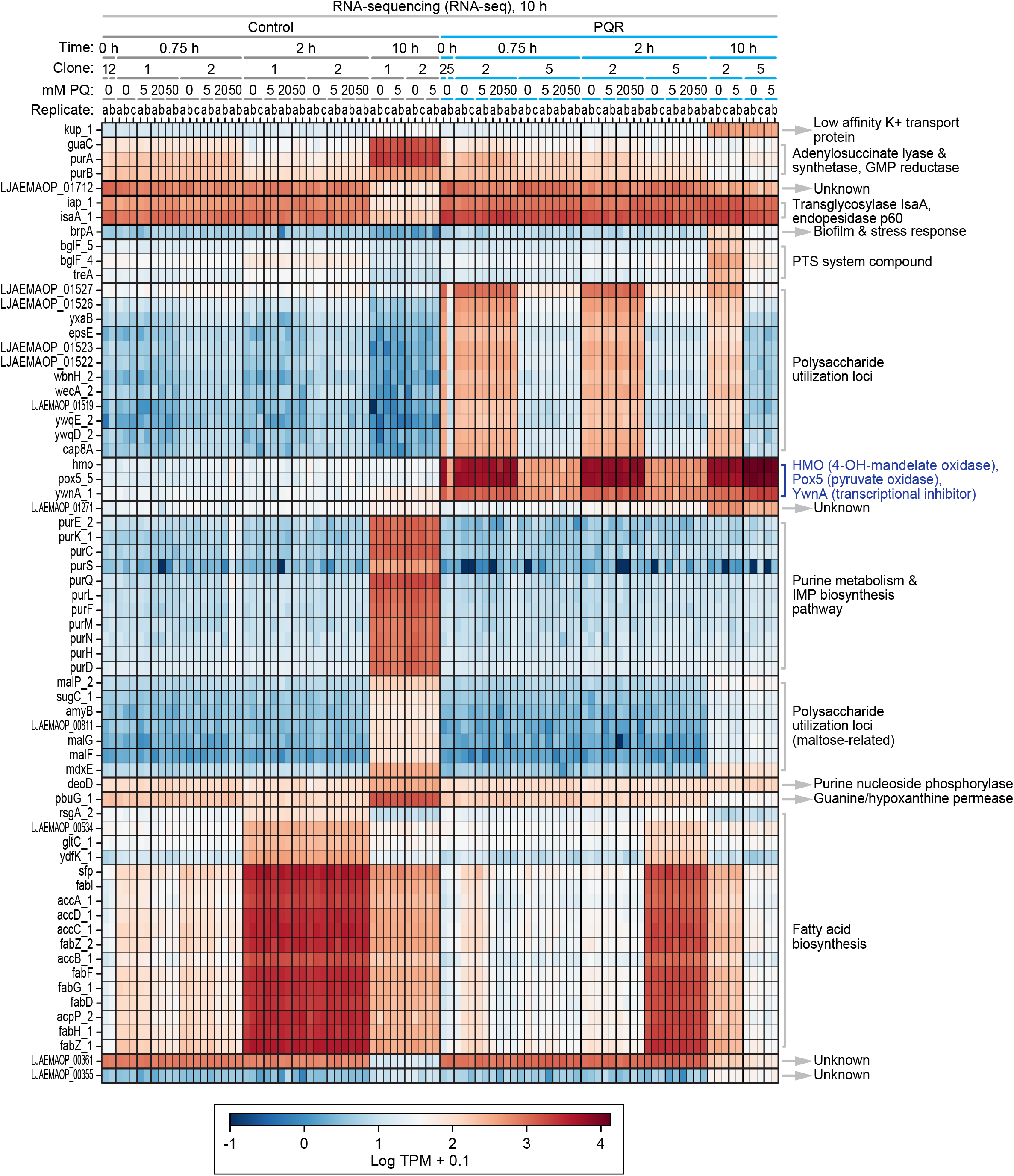
RNA-seq data revealed numerous changes in gene expression between PQR and control lines of *L. plantarum* regardless of short term paraquat treatment. Two isolates each of control *L. plantarum* (clones 1, 2) and PQR *L. plantarum* (clones 2, 5) were incubated in media with 0, 5, 20, or 50 mM paraquat. Replicate samples (a, b, or c) were removed at 0, 0.75, 2, and 10 hours of incubation and processed for RNA seq analysis. Significantly differentially expressed genes are highlighted in the heatmap (see Log TPM + 0.1 look up table legend at bottom). Major differences include IMP biosynthesis, polysaccharide utilization, ATP productions and synthesis, and the operon controlling acetate production, *ywnA*-*pox5*-*hmo* (dark blue text). See Table S1 for statistical comparisons and population size.

**Figure S4.**
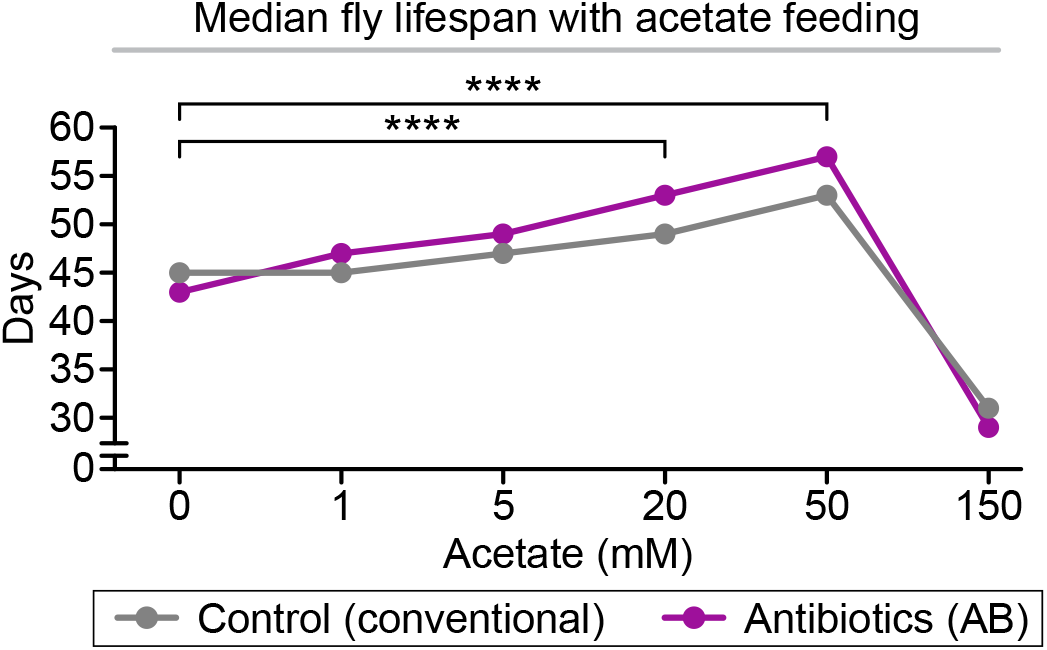
Feeding acetate to *Drosophila* extended lifespan in a dose-dependent manner, independent of an intact microbiome. Flies fed increasing doses of sodium acetate had increased lifespans up to 50 mM of concentration, even when also fed antibiotics. p-values were obtained by logrank analysis. See Table S1 for statistical comparisons and population size.

## METHODS

### *Ex vivo* directed evolution of *Drosophila melanogaster* gut microbiota

Whole Intestines from 15 conventionally reared adult *WCS* female flies were dissected in sterile PBS at 2 weeks post-eclosion. Intestines were homogenized in a microcentrifuge tube with a sterile plastic pestle. Intestinal bacteria were then cultured in MRS media (Sigma) overnight at 30°C. After 18 hours of growth, the bacterial culture was split into two separate culture conditions. One condition contained successive dilutions of paraquat including 0.1 mM, 0.5 mM, 1 mM, 5 mM, 10 mM, 25 mM, 50 mM, 100 mM, 150 mM, and 200 mM taken from a 1.5 M stock of paraquat dissolved in sterile distilled water. The same volume of water alone was used as a vehicle control for the other half of the split culture. 10 µLs of culture containing absorbance 0.1 at OD 600 nm were then used to populate each of the cultures and dilutions and bacteria were allowed to grow overnight (18 hours). The next morning, the cultures containing visible growth at the highest concentration of paraquat were then used to start new cultures of serially diluted paraquat in MRS. Initially, the bacteria could only grow in paraquat concentrations around 1-5 mM. After 25 passages in these gradually paraquat adapted cultures, growth of the bacteria could be regularly observed at paraquat concentrations of up to 150 mM. This population of bacteria was then utilized for subsequent experiments as the PQR population. The control population was bacteria passaged with only water vehicle for 25 times. Numerous glycerol stocks of each population were frozen at -80° C.

### Identification of bacterial species

Cultures of either PQR or Ctrl bacteria were diluted onto sterile MRS agar plates and grown overnight at 30° C. Thirty individual colonies were randomly selected from each condition, and replicate plated on MRS medium. These colonies were then sent out for universal 16S rDNA Sanger sequencing at GeneWiz (Azenta Life Sciences). Returned sequences were compared and identified using NCBI BLAST. Proportions of each bacterial species were quantified from these data. These individual isolates were then grown and frozen as glycerol stocks for downstream experiments.

### Fly strains

*w^11^*^18^ *Canton-S* (*WCS*) were used as the ‘wild-type’ strain throughout this paper. *esg-Gal4, UAS-GFP* flies were outcrossed 10 generations to the *WCS* background for analysis of mis-differentiated stem cells and Dihydroethidium (DHE) quantification. *Drosophila* were maintained at 25°C at 55-60% humidity in a 12-hour light-dark cycle. Female flies were utilized in all experiments except when otherwise noted.

### Fly media

Developmental media consisted of standard yeast-cornmeal-agar media (Archon Scientific, glucose recipe: 7.6% glucose, 3.8% yeast, 5.3% cornmeal, 0.6% agar, 0.5% propionic acid, 0.1% methyl paraben and 0.3% ethanol). Adult medium contained 3% yeast extract (Difco) 4% dextrose, 2% sucrose, 8% cornmeal, 1% agar, 0.5% propionic acid, 0.1% methyl paraben and 0.3% ethanol (all from Lab Scientific). For axenic media, the above recipes were used without agar, propionic acid, or methyl paraben and autoclaved. A solution of 10% agar was also made and autoclaved. The appropriate volume of 10% agar solution was added aseptically to the autoclaved recipes to 0.6% for developmental food or 1% for adult food and dispensed into sterilized vials or bottles. Bacterial repopulation medium contained sterilized liquid food with 4% dextrose, 2% sucrose, 3% yeast extract. For antibiotic (AB) medium and gut microorganism clearance, flies were placed on medium containing 500 μg/mL ampicillin, 50 μg/mL tetracycline and 200 μg/mL rifamycin in 50% ethanol (Sigma). For acetate feeding, sodium acetate (Sigma) was dissolved in sterile distilled water and added to the appropriate concentrations, water alone was used as the vehicle. All percentages are given in w/v except for propionic acid and ethanol which is given in v/v.

### Axenic fly rearing and maintenance

Embryos were collected by overnight ∼16-hour egg laying in sterile cages on medium containing 0.5% sucrose, 3% yeast extract, and 1% agar. Plates were rinsed in sterile PBS to remove embryos. Embryos in PBS were transferred to a 15 mL Falcon tube and pelleted at 500x*g* for 5 minutes. Supernatant PBS was removed, and embryos were washed 3 times for 5 minutes in 5% bleach (Clorox), 50% ethanol (Sigma) for sterilization and dechorionization. This was followed by 3 washes in sterile PBS. ∼100 µLs of embryos were seeded onto bottles containing sterile developmental medium (Archon recipe). Bottles were placed in sterilized Tupperware containers in the incubator with minimal openings to allow for gas exchange. Upon eclosion, flies were transferred to sterile adult medium, allowed to mate for 48 hours, and then sorted into groups of males and females on a sterilized fly pad, under light CO_2_ pressure (no more than 3 minutes of anesthetization). Sterility was monitored by sampling random vials of flies and performing Colony Forming Unit (CFU) assays, or 16S bacterial rDNA PCR. After another 48 hours, flies were then repopulated with appropriate bacterial culture. Fly vials were kept in sterilized plastic containers within the incubator.

### Microbial repopulation

Large overnight cultures (200 mL) of PQR or control bacterial populations were grown in MRS at 30°C. Bacteria were pelleted at 5000x*g* for 10 minutes. MRS supernatant was removed. Bacteria were then resuspended in liquid repopulation media (above) to a concentration of OD 0.3 at 600 nm. Sterile vials containing two sterilized Kimwipes packed at the bottom were then soaked in 3 mL of bacteria containing repopulation medium. Young (4 day old) females were collected and placed on the appropriate bacteria containing medium vials (25-30 flies/vial) and allowed to eat the bacterial repopulation medium for 3 days within the incubator. These, now bacterially repopulated flies were then transferred to axenic adult medium for further assays. For experiments where the bacteria were used as probiotics, conventionally reared (non-axenic) flies were fed repopulation medium containing the appropriate bacteria for 3 days every 10 days until day 45 of adulthood.

### Paraquat survival

Repopulated flies were fed sterilized adult medium containing 20 mM paraquat. Their survival was monitored twice each day. A minimum of 100 flies were used for each survival trial. Logrank analysis was used to compare survival curves.

### Lifespan analysis

*Drosophila* were reared from embryos in low-density bottles with either yeast-cornmeal-agar axenic food or glucose recipe media described above (Archon Scientific). Newly eclosed flies (approximately 24 h post-eclosion) were collected onto adult axenic medium and allowed to mate for 48 h. Female flies were then separated, microbially repopulated, and maintained at a density of 30–35 flies per vial in a humidified (60%), temperature-controlled (25°C) incubator with a 12-h light–dark cycle. Flies were flipped to fresh medium every two days and death was scored at time of flipping. Lifespans were compared by logrank analysis.

### Colony Forming Unit (CFU) Assay

10 flies per biological replicate of various conditions and microbial repopulation status were washed twice in 70% ethanol and twice in sterile PBS. Flies were then ground up by pestle in a microcentrifuge tube containing 500 µLs of sterile MRS. Serial dilutions of fly extracts were plated on MRS agar plates and allowed to grow overnight at 30°C. Colonies present at specific dilution were then counted, and CFUs per fly were calculated with the appropriate dilution factor. ANOVA followed by Tukey’s pot-hoc test were used to compare samples.

### Microbial load quantification by PCR

For 16S bacterial rDNA quantification, 4 replicates of 10 whole flies (washed twice in 70% ethanol and twice in PBS) were used for total DNA extraction via the Power Soil DNA isolation kit (Qiagen). Universal primers for the 16S ribosomal RNA gene were against variable regions 1 (V1F) and 2 (V2R).^8^

### Primers

*16Sr*-fwd-AGAGTTTGATCCTGGCTCAG

*16Sr*-rev-CTGCTGCCTYCCGTA

*Dm Actin5C*-fwd-TTGTCTGGGCAAGAGGATCAG

*Dm Actin5C*-rev-ACCACTCGCACTTGCACTTTC

### CAFE Assay

Analysis of capillary feeding (“the CAFE assay”) was performed similarly to (Ja *et. al* 2007) with minor modifications.^43^ Briefly, 10 flies were placed in vials with wet tissue paper as a water source and a capillary food source (3.8% glucose, 1.9% sucrose, 3% yeast extract, and 0.2% FD&C Blue No. 1) topped with mineral oil to prevent dehydration. Feeding was monitored for at least 24 hours, replacing depleted capillaries as necessary. A minimum of 8 groups of 10 flies per condition were used for each experiment.

### Dihydroethidium (DHE) staining and intestinal stem cell (ISC) and enteroblast (EB) quantification

Flies at 45 days of age were anesthetized on ice and intestines were dissected in cold Schneider’s Medium (Thermo Fisher Scientific). For ROS staining, dissected intestines were immersed in 50 µM DHE (Invitrogen) in Schneider’s Medium for 5 minutes at room temp, then washed three times for 30 seconds in Schneider’s Medium. DHE samples were quickly mounted in Schneider’s Medium and cover slips sealed with nail polish. Intestines were imaged for no more than 30 minutes on a Zeiss LSM800 confocal microscope with a 60x 1.4 N.A. oil immersion objective. Z-stacks spanning the entire posterior midgut were taken. Quantification of DHE was performed using FIJI,^44^ in which mean fluorescent intensity values were quantified. Quantification of area and number of ISCs/EBs was determined in FIJI. A minimum of 9 midguts were used for each treatment. Comparisons were made by a two-tailed Student’s t-test.

### Intestinal barrier dysfunction “Smurf” assay

The “Smurf” fly intestinal barrier dysfunction assay was performed as previously described.^12,19^ Axenic flies were repopulated with PQR and control microbiota and aged on standard **adult medium** (see ‘Fly media’) until the day of the Smurf assay. Dyed medium was prepared by the addition of FD&C Blue No. 1 at a final concentration of 2% w/v. A fly was counted as a Smurf when dye coloration was observed outside the digestive tract. Comparisons of Smurf proportion per time point were carried out using binomial tests to calculate the probability of having as many Smurfs in population A as in population B, as well as analysis of variance (ANOVA) for proportions of Smurf flies per replicate vial with a minimum of 7 vials of 12–31 flies per replicate.

### Statistical analysis

We assessed the normality of our data using the D’Agostino-Pearson omnibus normality test and the Shapiro-Wilk normality test, which have good power properties over a wide range of distributions.^45^ For datasets that passed both normality tests, we used the unpaired Student’s t-test with Welch’s correction when comparing two groups and the one-way ANOVA (Dunn’s multiple comparisons) when comparing three or more groups. For data sets that failed either one of the normality tests, we used the Mann-Whitney U test when comparing two groups and the Kruskal-Wallis test with Dunn’s post hoc test when comparing three or more groups. Lifespan and survival curves were assessed using logrank analysis. All statistical analyses were performed using GraphPad Prism. Significance was expressed as p-values (n.s.=p>0.05, *=p≤0.05, **=p≤0.01, ***=p≤0.001, ****=p≤0.0001).

### Bacteria RNA-seq

#### RNA extraction

Bacterial RNA was extracted using RNAsnap methods.^46^ Briefly, frozen bacterial pellets were suspended in 500 µL RNAsnap mix (95% formamide, 18 mM ethylenediaminetetraacetic acid, 0.025% sodium dodecyl sulphate, 1% β-mercaptoethanol) before addition of ∼200 µL of 0.1 mm zirconia silica beads (Biospec 11079101Z). Cells were then lysed by bead beating for 3 × 2.5 minutes in a Biospec Mini Bead Beater (Biospec 1001) with 5-minute intervals and then subjected to centrifugation at 4,300x*g* for 5 minutes. The clean supernatant was then transferred to a new tube, and RNA was purified using a Zymo ZR-96 RNA Clean & Concentrator Kit (Zymo R1080) per manufacturer’s instructions.

#### Library preparation

RNA-seq libraries were constructed following a modified RNA tag-seq protocol as detailed in (Huang et al. 2020, *Nucleic Acids Research*).^47^ Briefly, 400 ng of total RNA lysate was subjected to fragmentation in 2× FastAP buffer (ThermoFisher EF0651), genomic DNA removal (TURBO DNase, ThermoFisher AM2239) and dephosphorylation (FastAP, ThermoFisher EF0651). Fragmented RNA was purified using SeraPure SPRI bead cleanup and ligated with barcoded first adapter ligation by T4 RNA ligase I (NEB M0437M). After pooling and purification with the Zymo RNA Clean & Concentrator-5 Kit (Zymo R1015), we quantified barcoded RNA using a Qubit RNA HS Assay Kit (ThermoFisher Q32855) and then performed RNase-H-based ribosomal RNA depletion on 400 ng of barcoded RNA sample using a 10:1 probe-to-RNA ratio. rRNA-depleted RNA was subjected to downstream library preparation following standard RNAtag-seq protocols, including reverse transcription (ThermoFisher 18090010) and second adapter ligation (NEB M0437M). Ligation products were further amplified with primers containing Illumina P5 and P7 adapters and sample indexes, and polymerase chain reactions (PCRs) were stopped during exponential amplification. PCR products were subjected to gel electrophoresis on E-Gel EX Agarose Gels, 2% (ThermoFisher G402002) and expected DNA smears (300–600 bp) were excised and extracted using the Monarch DNA Gel Extraction Kit (NEB T1020L). Sequences of all adapters and primers used in library preparation are provided in reference.^47^

#### Data analysis

RNA libraries were analyzed for differential expression analysis as outlined in (Huang et al. 2020, *Nucleic Acids Research*)^47^, which lists all adapter and primer sequences used. Briefly, raw sequencing reads were demultiplexed using Sabre and bcl2fastq before adapter removal with Cutadapt v2.1, using parameters ‘-a file:[RNAtagSeq adapter.fa] -u 5 –minimum-length 20 –max-n 0 -q 20’ to remove low-quality bases and adapters. To mitigate the effect of rRNA reads, we performed alignments against the 16S rRNAs of corresponding strains with Bowtie2, using the versions and parameters outlined in reference.^47^ Genomes used for sequencing alignments were de novo assembled as described in reference.^47^ The number of reads uniquely mapped to each coding sequence was calculated using featureCounts v1.6.2 without restraint on strandness (-s 0), and the expression level of each coding sequence (CDS) by transcripts per million (TPM) was quantified using an in-house script.

### Whole genome sequencing and SNP analysis

#### gDNA extraction

40 µL 0.1 mm Zirconia Silica beads (Biospec, 11079101Z) and 120 µL lysis solution (50 mM Tris–HCl, pH 7.5 and 0.2 mM EDTA) were added to a bead beating tube. Next, 40 µL bacteria culture were added and the tubes were centrifuged for 1 minute at 4,500x*g*. Then, the tubes were fixed on a bead beater (Biospec, 1001) and subjected to bead beating for 5 minutes, followed by a 10-min cooling period. The bead beating cycle was repeated once and tubes were centrifuged at 4,500x*g* for 5 min to spin down cell debris. Next, 10 µL cell lysate was transferred to a PCR tube (Bio-Rad, HSP3801) and 2 µL proteinase K solution (50 mM Tris–HCl, pH 7.5 and 1 µg/µL^−1^ proteinase K (Lucigen, MPRK092)) was added. Finally, cell lysate was subjected to proteinase K digestion on a thermal cycler (65°C 30 minutes, 95°C 30 minutes, 4°C infinite) and transferred the next day to −20°C for long-term storage.

#### Whole genome sequencing

Paired-end libraries were constructed following a published protocol of low-volume Nextera library preparation^48^ and sequenced on Illumina Nextseq 500/550 platform (2 × 75 bp) and HiSeq platform (2 × 150 bp). Raw reads were then processed by Cutadapt v2.1 with the following parameters: ‘--minimum-length 25:25 -u 10 -u -5 -U 10 -U -5 -q 15 --max-n 0 --pair-filter=any’ to remove low-quality bases and Nextera adapters. Coverage was 1.42 ± 2.86 million paired-end reads per isolate and PacBio long-read sequencing was performed for some isolates by SNPsaurus to improve the performance of *de novo* genome assembling.

#### Data analysis

Illumina reads passing quality filtering and PacBio long reads were assembled by Unicycler v0.4.4 with default setting to generate draft genomes, and the quality and species-level taxonomy of draft genomes were then assessed by QUAST v4.6.3, CheckM v1.0.13 and GTDB-Tk v0.2.2. Illumina reads of isolates were aligned to reference genomes by Bowtie2 v2.3.4 in paired-end mode with ‘--very-sensitive’ setting. Resulting reads alignments were then processed by SAMtools v1.9 and BCFtools v1.9 with ‘--ploidy 1’ setting to call genomic variation (SNPs and Indels).

#### Additional 16S rRNA sequencing

The same gDNA was used for 16S rRNA sequencing. The variable regions 3 and 4 (V3–V4) of the 16S rRNA gene was amplified by PCR using dual-index primers and sequenced on Illumina paired-end platform to generate 250 bp paired-end raw reads. For the 16S rRNA sequencing data analysis, raw sequencing reads of 16S-V4 amplicon were analyzed by USEARCH v11.0.667. Specifically, paired-end reads were merged using “-fastq_mergepairs” mode with default setting. Merged reads were then subjected to quality filtering using “-fastq_filter” mode with the option “-fastq_maxee 1.0 -fastq_minlen 240.” Remaining reads were deduplicated (-fastx_uniques) and clustered into OTUs (-cluster_otus) at 97% identity with the option “-minsize 2”, and merged reads were then searched against OTU sequences (-otutab) to generate OTU count table. Taxonomy of OTUs were assigned using Ribosomal Database Project classifier v2.13 trained with 16S rRNA training set 18. Relative abundances were defined as read counts of specific OTU normalized by total number of non-spike-in reads, and absolute abundances were defined as reads count of specific OTU normalized by weight of input and reads count of spike-in strain OTU.

#### Acetate quantification

Monocultures of *Lactiplantibacillus plantarum* (Control or PQR) were allowed to grow to stationary phase for 18 hours in MRS medium. Bacteria were pelleted at 10,000x*g* for 10 minutes. Supernatant from the media of individual cultures was taken and used in the acetate quantification colorimetric enzymatic assay (Bio Assay Systems). A standard curve of sodium acetate was used to generate a linear signal range of absorbance at 570 nM. Cultured supernatant was added at a concentration sufficient to fall within the standard curve. Sterile MRS media alone was used as the background signal control for cultured acetate production. A minimum of four cultures per condition were used to determine acetate production into the media and repeated 3 times. A two-tailed Student’s t-test was used to compare acetate levels in control and PQR medium.

## AUTHOR CONTRIBUTIONS

MU and MSH conceived the experiments. Experiments were performed and analyzed by MU (husbandry, lifespan, intestinal staining, bacterial load, barrier function, secondary data analysis, acetate feeding lifespans and acetate quantification), YH and YS (RNA/DNA-sequencing secondary data analysis), TC (heat maps), CL (microbial repopulation). MU, JCC, HW, YW, YS, TC and MSH made intellectual contributions, designed the figures, and wrote the manuscript.

## ACKNOWLEDGMENTS

We thank all members of the Ulgherait, Wang, Canman, and Shirasu-Hiza labs for support, discussions, and feedback. Support for this research came from the AFAR Glenn Foundation Postdoctoral Fellowship for Aging Research (MU), NSF GRFP (TC, CL), NIH R01GM117407 (JCC), NIH R01GM130764 (JCC), NIH 2R01AI132403 (HHW), NIH 1R01EB031935 (HHW), NIH 1R01DK118044 (HHW), NSF2522218 (HHW), NIH R35GM127049 (MSH), and NIH R01AG045842 (MSH).

## DATA AVAILABILITY

The authors declare that all data supporting the findings of this study are available, including replicate experiments, and will be made available upon reasonable request to the first author, Matt Ulgherait.

